# Rapid evolution of the embryonically-expressed homeobox gene *LEUTX* within primates

**DOI:** 10.1101/2023.02.02.526829

**Authors:** Thomas D. Lewin, Josephine R. Blagrove, Peter W. H. Holland

## Abstract

LEUTX is a homeodomain transcription factor expressed in the very early embryo with a function around embryonic genome activation. The *LEUTX* gene is found only in eutherian mammals, including humans, but unlike the majority of homeobox genes, the encoded amino acid sequence is very different between divergent mammalian species. However, whether dynamic evolution has also occurred between closely related mammalian species remains unclear. In this work, we perform a comparative genomics study of *LEUTX* within the primates, revealing dramatic evolutionary sequence change between closely related species. Positive selection has acted on sites in the LEUTX protein, including six sites within the homeodomain; this suggests that selection may have driven changes in the set of downstream targets. Transfection into cell culture followed by transcriptomic analysis reveals small functional differences between human and marmoset LEUTX, suggesting rapid sequence evolution has fine-tuned the role of this homeodomain protein within the primates.

**Significance:** Homeobox genes are key regulators of animal development and are therefore often highly conserved between divergent species. However, recent work has uncovered several apparently fast-evolving homeobox genes expressed during mammalian development, one of which is *LEUTX*. Here we show that the *LEUTX* genes of primates are highly variable, and that these sequence changes have resulted in small modifications to its protein function. This continued rapid divergence of an embryonic gene between closely related species is significant to the areas of evolutionary, developmental and genome biology.

## Introduction

Homeobox genes are renowned for their conservation across large evolutionary timescales. Many homeodomain (HD) transcription factors (TFs) play essential roles in fundamental animal developmental processes, such as axial patterning, cellular differentiation and cell proliferation (Duboule 1994; Gehring et al. 1994; Bürglin & Affolter 2016), and have been at the centre of the idea of the conserved genetic toolkit due to their striking similarity across widely divergent animal species (Carroll 2000, 2008).

It is therefore intriguing that in recent years an increasing number of homeobox genes have been found to be lineage-specific and rapidly evolving, contrary to the evolutionary conservation typical of this group. This is particularly the case for the PRD-like class of homeobox genes, to which the fast-evolving and mammal-specific Cphx, Dux, Rhox, and Eutherian Totipotent Cell Homeobox (ETCHbox) genes belong. All of these sets of genes have been recruited to roles in early mammalian development (Leidenroth & Hewitt 2010; Hendrickson et al. 2017; Li et al. 2006; Madissoon et al. 2016; Maeso et al. 2016; MacLean & Wilkinson 2010).

The ETCHbox genes duplicated from the *CRX* homeobox gene in the ancestor of eutherians and the last eutherian common ancestor possessed six group members: *ARGFX*, *DPRX*, *LEUTX*, *PARGFX*, *TPRX1* and *TPRX2* (Maeso et al. 2016). ETCHbox genes are expressed exclusively during early preimplantation development (Maeso et al. 2016); recent work has shown that they function around or immediately after embryonic genome activation (EGA) in humans and mice, with early transcriptional programs defective when they are knocked down (Zou et al. 2022; Guo et al. 2022; Jouhilahti et al. 2016; Madissoon et al. 2016; Töhönen et al. 2015). Mouse ETCHbox genes are necessary for proper blastocyst development and hatching (Cui et al. 2016), and we recently showed that bovine ETCHbox genes have probable roles in blastocyst formation (Lewin et al. 2022). Moreover, the ETCHbox gene *TPRX1* is necessary for transforming pluripotent human embryonic stem cell cultures into totipotent 8-cell-like cells, suggesting a role in totipotency (Mazid et al. 2022).

This body of work implies that ETCHbox proteins are critical regulators of developmental processes in the mammalian preimplantation embryo. The paradox is that, despite these roles, ETCHbox genes seem to be evolving rapidly. ETCHbox repertoires have undergone dramatic evolutionary changes across the eutherians, with high rates of both gene duplications and losses leading to dramatic copy number variation between species (Lewin et al. 2021; Katayama et al. 2018; Maeso et al. 2016). An illuminating comparison is between human, which has lost just *PARGFX* and has a single copy of the other five genes, and mouse, in which *ARGFX*, *DPRX*, *PARGFX* and *LEUTX* are all lost or pseudogenised, *TPRX1* is present in two copies and *TPRX2* in 66 copies (Maeso et al. 2016; Royall et al. 2018). Other large tandem arrays of ETCHbox genes have been found in *Oryctolagus cuniculus* (rabbit – 27 *LEUTX* copies) and *Myotis myotis* (greater mouse-eared bat – six *TPRX2* copies) (Lewin et al. 2021). However, previous work employed a broad sampling strategy, leaving open the question of whether closely related species possess different ETCHbox repertoires.

The differences between mammalian taxa are not restricted to gene duplication and loss. ETCHbox genes and their ‘ancestor’ *CRX* exhibit asymmetric sequence evolution: *CRX* has been conserved while ETCHbox sequences have diverged extensively between taxa, and this divergence has been driven at least in part by positive selection (Maeso et al. 2016; Lewin et al. 2021). In previous work, we compared the transcriptional activity of ETCHbox genes between humans, mice and cattle, and found evidence for changes in gene function (Lewin et al. 2022); we define ‘function’ here as the gene sets up- and down-regulated by a putative transcription factor.

Overall, previous work has shown that extensive sequence divergence and changes in ETCHbox protein function are seen between deeply diverged evolutionary lineages of eutherian mammals. We asked to what extent there have been changes between more closely related species. This will help answer whether ETCHbox homeobox genes are ‘fast-evolving’ or whether they underwent change during mammalian diversification followed by relative stasis. To address this question, we characterised the ETCHbox gene *LEUTX* across the order Primates, with species spanning from a few million years to circa 75 million years of divergence (Pozzi et al. 2014; Wilkinson et al. 2011; dos Reis et al. 2018). We find that *LEUTX* sequences have continued to diverge at a rapid rate across primates and that positive selection has driven substitutions at key HD residues, suggesting selection for divergence of protein function. Experimental characterisation using transfection followed by RNA-seq suggests small but significant differences exist in the TF function of LEUTX between primate species.

## Results

### Duplication and divergence of *LEUTX* within primates

We identified the *LEUTX* genes in publicly-available genome sequences of 52 primate species representing all major evolutionary lineages (Figure 1A, B; Supplementary Figure S1; Supplementary Table S1). Although *LEUTX* has been lost in several other mammals (Lewin et al. 2021), the gene is present in all of the sampled primate genomes. Of 52 species analysed, 48 have a single putatively functional *LEUTX* locus in the expected location in the genome. We find four species with duplications: (a) ten *LEUTX* loci in *Microcebus murinus* as reported previously (Lewin et al. 2021); (b) two tandem *LEUTX* loci in *Lemur catta*; (c) a divergent, intron-containing copy on a separate scaffold in *Nycticebus bengalensis*; (d) a partial gene duplication affecting exons 1 and 3 in *Hylobates pilateus*. The first three examples are all members of the Strepsirrhini, which includes the lemurs and lorises (Figure 2A).

**Fig. 1.**
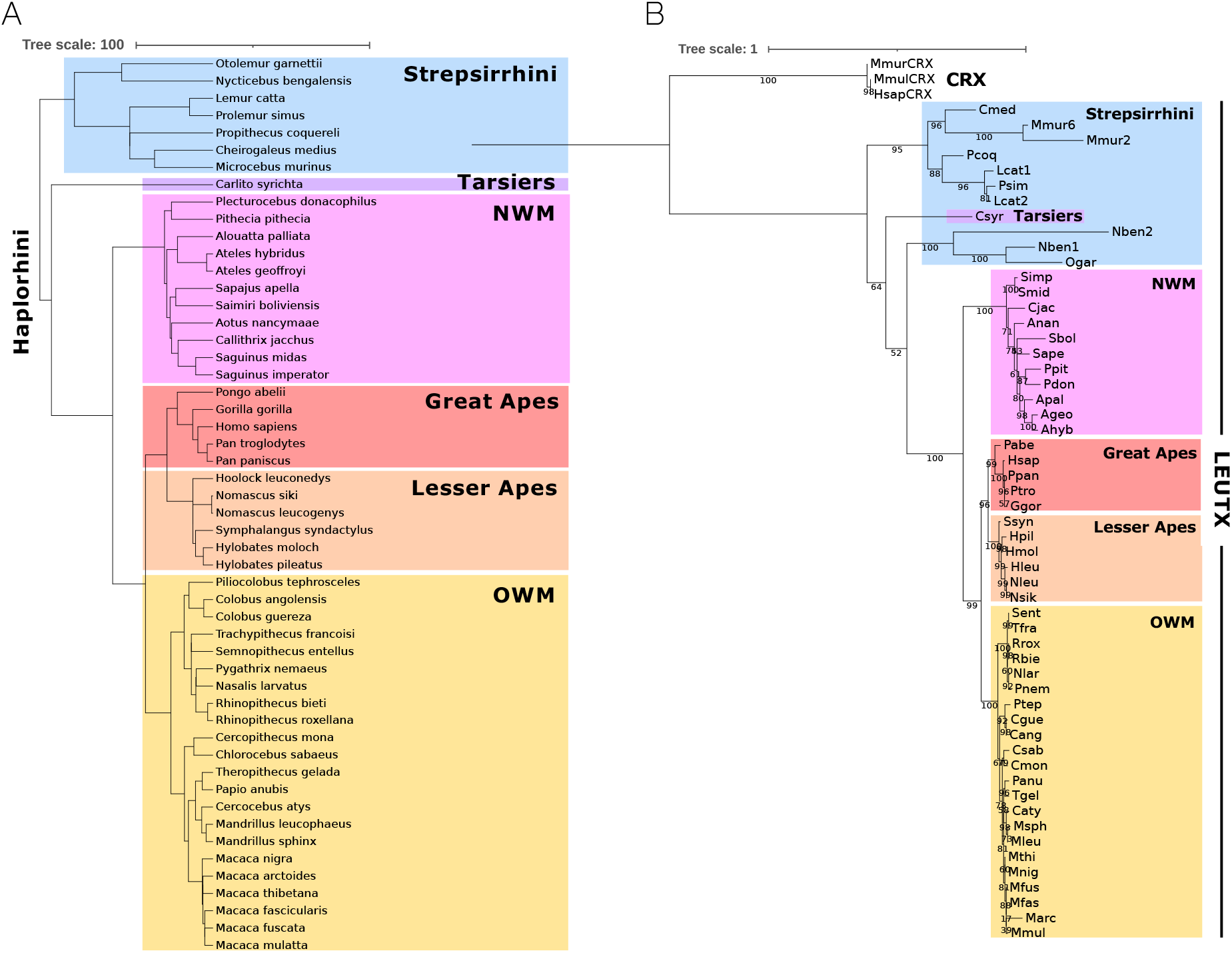
Phylogenetic analysis of primate LEUTX sequences. **A)** Species tree of the 52 primates used in this analysis. Branch lengths are proportional to divergence times (million years). **B)** Tree of primate LEUTX sequences made with full protein-coding sequences. Great ape, lesser ape, Old World monkey and New World monkey clades are recapitulated with the expected topology. Nben and Ogar are expected to group with the lemurs (Cmed, Mmur, Pcoq, Lcat, Psim). Abbreviations: Ageo = *Ateles geoffroyi*; Ahyb = *Ateles hybridus*; Anan = *Aotus nancymaae*; Apal = *Alouatta palliate*; Cang = *Colobus angolensis*; Caty = *Cercocebus atys*; Cgue = *Colobus guereza*; Cimi = *Cebus imitator*; Cjac = *Callithrix jacchus*; Cmed = *Cheirogaleus medius*; Cmon = *Cercopithecus mona*; Csab = *Chlorocebus sabaeus*; Csyr = *Carlito syrichta*; Ggor = *Gorilla gorilla*, Hleu = *Hoolock leuconedys*; Hmol = *Hylobates moloch*; Hpil = *Hylobates pileatus*; Hsap = *Homo sapiens*; Lcat = *Lemur catta*; Marc = *Macaca arctoides*; Mfasc = *Macaca fascicularis*; Mfus = *Macaca fuscata*; Mleu = *Mandrillus leucophaeus*; Mmul = *Macaca mulatta*; Mmur = *Microcebus murinus*; Mnig = *Macaca nigra*; Msph = *Mandrillus sphinx*; Mthi = *Macaca thibetana*; Nben = *Nycticebus bengalensis*; Nlar = *Nasalis larvatus*; Nleu = *Nomascus leucogenys*; Nsik = *Nomascus siki*; NWM = New World Monkeys; Ogar = *Otolemur garnettii*; OWM = Old World Monkeys; Pabe = *Pongo abelii*; Panu = *Papio anubis*; Pcoq = *Propithecus coquereli*; Pdon = *Plecturocebus donacophilus*; Pnem = *Pygathrix nemaeus*; Ppan = *Pan paniscus*; Ppit = *Pithecia pithecia*; Psin = *Prolemur simus*; Ptep = *Piliocolobus tephrosceles*; Ptro = *Pan troglodytes*; Rbie = *Rhinopithecus bieti*, Rrox = *Rhinopithecus roxellana*; Sape = *Sapajus apella*; Sbol = *Saimiri boliviensis*; Sent = *Semnopithecus entellus*; Simp = *Saguinus imperator*; Smid = *Saguinus midas*; Ssyn = *Symphalangus syndactylus*; Tfra = *Trachypithecus francoisi*; Tgel = *Theropithecus gelada*.

**Fig. 2.**
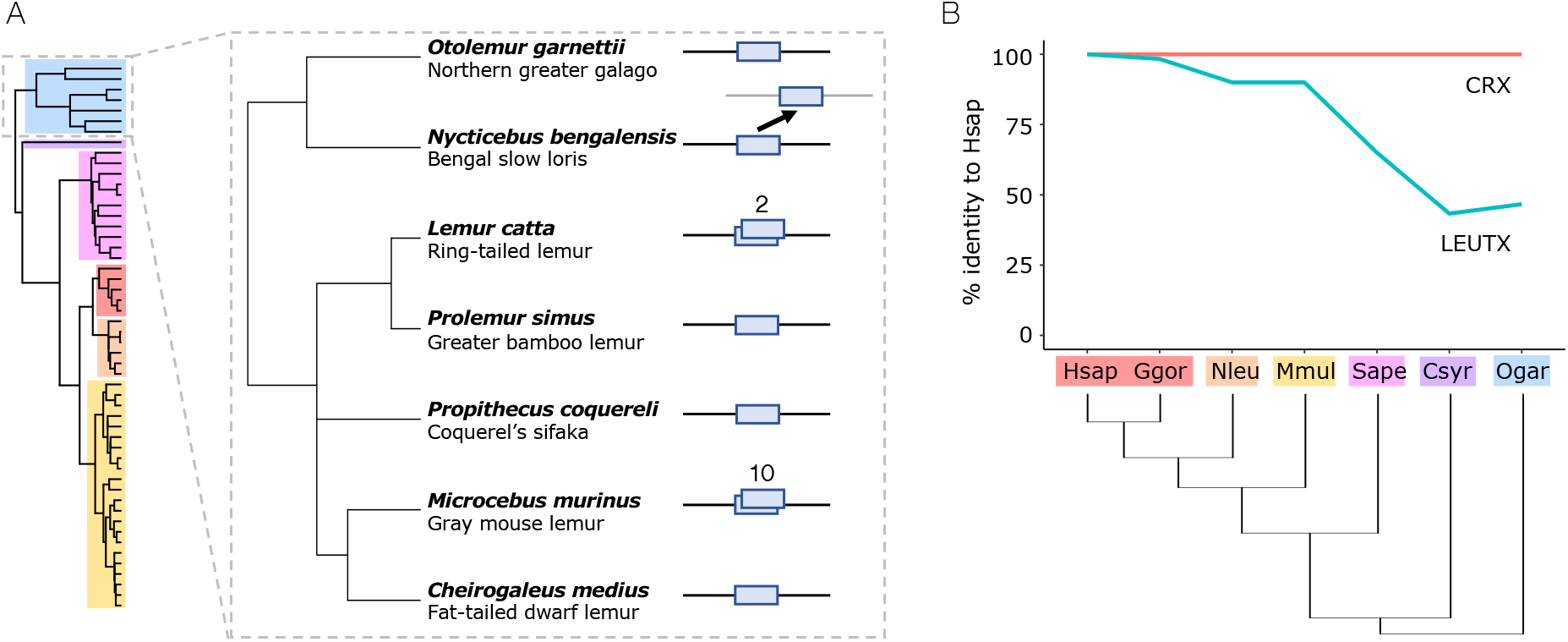
Evolution of LEUTX within primates. **A)** *LEUTX* copy number within strepsirrhine primates. *Nycticebus bengalensis* has a divergent second *LEUTX* copy on a separate scaffold. *Lemur catta* has two *LEUTX* tandem duplicates, and *Microcebus murinus* has 10 *LEUTX* loci. **B)** Divergence in primate LEUTX HDs. Plot shows percent identity of representative species’ LEUTX HDs to that of human. CRX is shown for reference. Abbreviations as in Figure 1.

We asked whether LEUTX protein sequences are fast-evolving within the primates. Using all vs. all pairwise comparisons of sequence identity, we find that primate LEUTX HDs show extensive divergence, increasing gradually with phylogenetic distance (Figure 2B). The two most different LEUTX HDs (*C. jacchus and M. murinus*) share just 35% sequence identity. Indeed, across the full coding sequence, only 12% (23/198) of amino acid sites are invariable between all sampled primates. Coding sequences are most variable within the Strepsirrhini; Figure 1B and Supplementary Table S2). This contrasts markedly with *CRX*, from which *LEUTX* originated by gene duplication, which is highly conserved across species as typical for homeobox genes (Figure 2B). Across a sample of 20 species representing all major evolutionary lineages (Supplementary Figure S2), 19 of the CRX HDs are identical, while *Pan troglodytes* has one substitution (A18T). Overall, we show that *LEUTX* protein-coding sequences have evolved rapidly within the primate lineage, including within the HD.

The primate CRX sequences show no variation in gene structure: the start and stop codons and intron/exon boundaries are conserved, and there are no indels. In contrast, of the 52 LEUTX sequences analysed, there are six different predicted start codons and seven different stop codon positions. For example, there are different predicted start codons in: Old World monkeys/apes (x2); New World monkeys (x2); tarsiers; and lemurs. Additionally, we uncover indels at four separate locations (Supplementary Figure S3). Overall, within primates, we observe *LEUTX* duplication, rapid sequence evolution and significant changes to gene structure, but no cases of *LEUTX* gene loss.

Within the genus *Macaca*, we were able to test the extent of variation between very closely related species. Among six species, we find two have identical deduced LEUTX proteins (*M. fascicularis* and *M. fuscata*), two differ from this reference by one substitution (*M. thibetana* and *M. nigra*; both H101R); and one species has a different substitution (*M. mulatta*; P92S) (Supplementary Figure S4). However, *M. arctoides* has 11 amino acid differences: six of these due to a frameshift-causing indel 16 residues from the end of exon 3, changing the last seven amino acids of the protein and creating a premature stop codon. We find there is more difference between LEUTX protein sequences within the *Macaca* genus than there is between CRX protein sequences across the entire primate order.

### Evolution of functional motifs

We tested whether positive selection has been a driver of LEUTX sequence divergence. We detected evidence for episodic diversifying selection within the primate lineage using the branch-site model of BUSTED (Murrell et al. 2015) (likelihood ratio test [LRT] *p*-value = 9.322 × 10^−7^). Analysis using MEME (Murrell et al. 2012) indicated that 27 residues within the protein have been under positive selection at some point in the primate phylogeny, six of which lie within the HD (Figure 3A, Supplementary Figure S3). This suggests that positive selection has played a role in the divergence of LEUTX proteins.

**Fig. 3.**
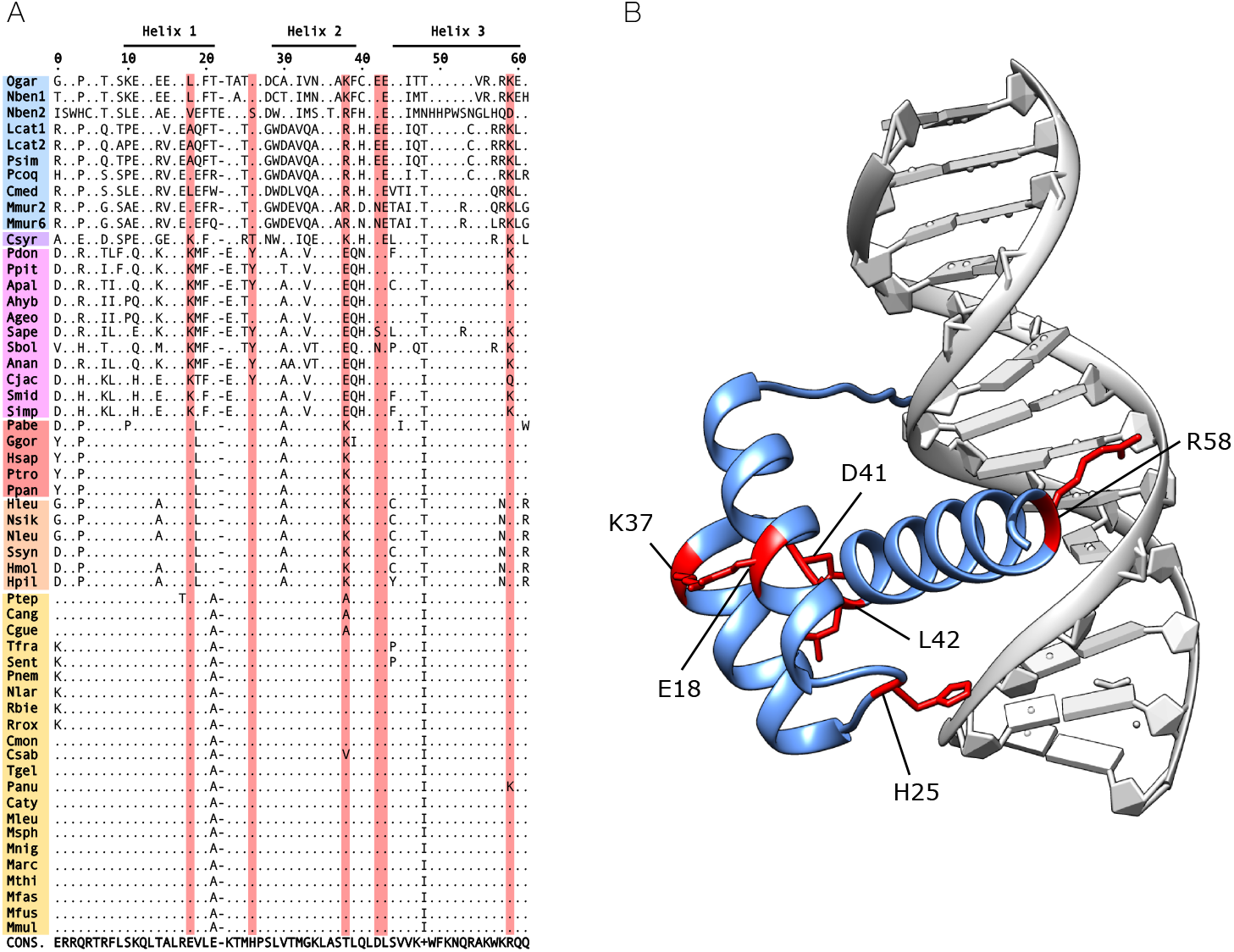
Positive selection in LEUTX homeodomains. **A)** Primate LEUTX homeodomains. Only residues divergent from the consensus are shown; consensus sequence is below alignment. Residues under positive selection are highlighted in red. Abbreviations as in Figure 1. **B)** Structure of human LEUTX homeodomain (blue) in complex with DNA (grey). Residues in which positive selection was detected in primates are shown in red; side chains shown only for these residues.

One of the residues inferred to have changed due to positive selection is HD residue 58, found within the critical ‘recognition helix’ (helix 3), which the structural modelling approach of Katayama et al. (2018) identified as a specificity-determining residue in LEUTX. In almost all Old World monkeys and apes (including human) this residue is R58, while the majority of New World monkeys and prosimians (strepsirrhines plus tarsiers) have K58. *C. jacchus* is notable for its unusual Q58 residue. Comparative structural modelling suggests that residue 58 contacts the major groove of the DNA double helix (Figure 3B). This suggests that within primates there has been selection for divergent specificity of LEUTX DNA-binding properties. Modelling also suggests that the side chains of residues under positive selection in HD helices 1 and 2, at positions 18 and 37 respectively, are in close proximity. Their opposite charges and HD position (Clarke et al. 1994) suggest the formation of salt bridges between these residues, implying selection for possible changes in the structure or stabilisation of the HD. Positively selected residue H25 contacts the DNA sugar-phosphate backbone.

Changes to other specificity-determining residues, as defined by Katayama et al. (2018), have also occurred, but are not confirmed as under positive selection with the current data set. First, A54 to V54 in *O. garnettii* and *N. bengalensis* (galago and loris). Second, position 47 has ‘flipped’ between I47 and T47 several times: T47 is seen in prosimians and New World monkeys, changing to I47 in *C. jacchus*; I47 is also seen in Old World monkeys, but changed to T47 in the ancestor of apes, again reverting to I47 in *Homo*, *Pan* and *Gorilla*. This complex evolutionary history suggests lability in this part of the LEUTX protein, consistent with previous work, which found this site has minimal functional influence alone but may undergo compensatory substitutions in response to changes at position 54 (Katayama et al. 2018). Other known specificity-determining residues (R2, R3, R5, K50, and N51) are invariant across primates (with the exception of the divergent *N. bengalensis* duplicate), and we identify pervasive purifying selection at R2, R5 and N51, along with 11 other residues within the HD (Supplementary Figure S3).

Katayama et. al (2018) annotated a ‘Leutx domain’, a conserved peptide motif downstream of the HD with conservation across mammals. Within this region, the authors propose two 9-amino acid transactivation domains (9aaTADs) in every mammalian sequence analysed; 9aaTADs are known to mediate the activation of transcription and are therefore key to TF function (Piskacek et al. 2007). We find both 9aaTADs are highly conserved across 44 anthropoids analysed (New World monkeys, Old World monkeys and apes); we detect evidence for purifying selection at four residues in the first 9aaTAD and two in the second (Supplementary Figure S3). As above, increased change is observed within the prosimians.

We also asked whether ubiquitination motifs in LEUTX showed evolutionary conservation across primates. Using an evolutionary screening algorithm (Jyun-Rong Wang et al. 2017), we identified three high-likelihood putative ubiquitination motifs in human LEUTX (Supplementary Figure S3). Each is conserved across all anthropoids, suggesting evolutionary constraint, but prosimians show notable divergence. For instance, *O. garnettii* is missing the target lysine at two out of three motifs, but putatively compensatory lysine substitutions are present within both of these motifs. Several other species are missing the target lysine in the ubiquitination motifs but evolved new lysine residues elsewhere. The conservation of ubiquitination motifs across anthropoids and the evolution of putative compensatory changes in prosimians points to functional importance, consistent with the genes’ fleeting temporal expression and subsequent requirement for rapid degradation.

### Evolution of LEUTX expression profiles

We asked whether the expression profiles of *LEUTX* in the preimplantation embryo are conserved across primates. Human *LEUTX* is expressed in a distinct temporal pattern, with expression peaking sharply at the 8-cell stage (Maeso et al. 2016). We quantified *LEUTX* expression across preimplantation development in publicly available human, *M. mulatta* (Old World monkey) and *C. jacchus* (New World monkey) RNA-seq datasets and found strong conservation of 8-cell stage-specific expression between human and *M. mulatta* (Figure 4). In *C. jacchus*, *LEUTX* is expressed in a more protracted pulse comprising both the 4-cell and 8-cell stages. Thus, *LEUTX* expression profiles can vary but remain constrained within the limits of embryonic genome activation and morula formation.

**Fig. 4.**
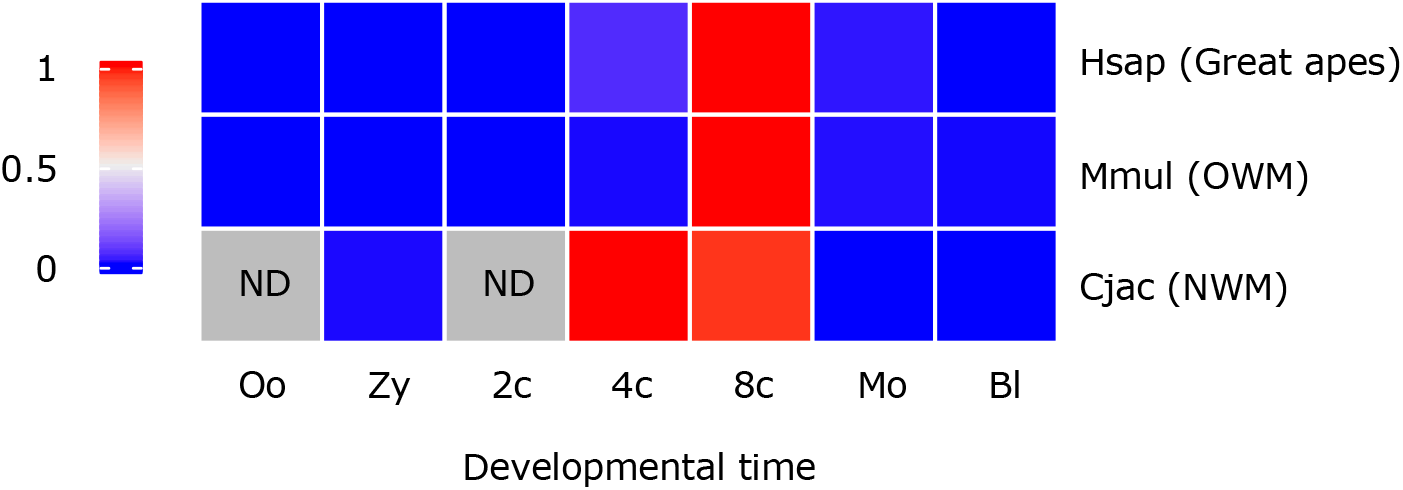
LEUTX expression in primate preimplantation embryos. Heatmap of scaled *LEUTX* expression in human (Hsap), rhesus macaque (Mmul) and common marmoset (Cjac) preimplantation embryos. Grey squares indicate no data available. Abbreviations: 2c = two-cell; 4c = four-cell; 8c = 8-cell; Bl = blastocyst; Cjac = *Callithrix jacchus*; Hsap = *Homo sapiens*; Mmul = *Macaca mulatta*; Mo = morula; ND = no data; NWM = New World monkeys; Oo = oocyte; OWM = Old World monkeys; Zy = zygote.

### Evolutionary divergence of LEUTX downstream targets

We hypothesised that the selection-driven sequence divergence observed between primates has caused divergence of LEUTX protein functions. We used transcriptome analysis after transfection into cultured cells to test this, targeting *Homo sapiens* (representing great apes) and the common marmoset *C. jacchus* (New World monkeys) for experimental comparison. The *C. jacchus* LEUTX HD has 73% sequence identity to human, including substitutions at four sites within the HD at which we identified positive selection, one of which is the specificity-determining residue 58 (Figure 5A). *LEUTX* gene sequences of *H. sapiens* and *C. jacchus*, each with a C-terminal V5 tag, were cloned into a constitutive mammalian expression vector and transfected into human dermal fibroblasts (HDFs). Previous work has shown that expression of ETCHbox genes in a cell culture setting, including in fibroblasts, elicits changes to the expression of embryonic genes (Maeso et al. 2016; Royall et al. 2018; Lewin et al. 2021; Madissoon et al. 2016; Jouhilahti et al. 2016). Immunocytochemistry confirmed protein expression and nuclear localisation of the homeodomain TF in both human and marmoset-transfected samples (Figure 5B).

**Fig. 5.**
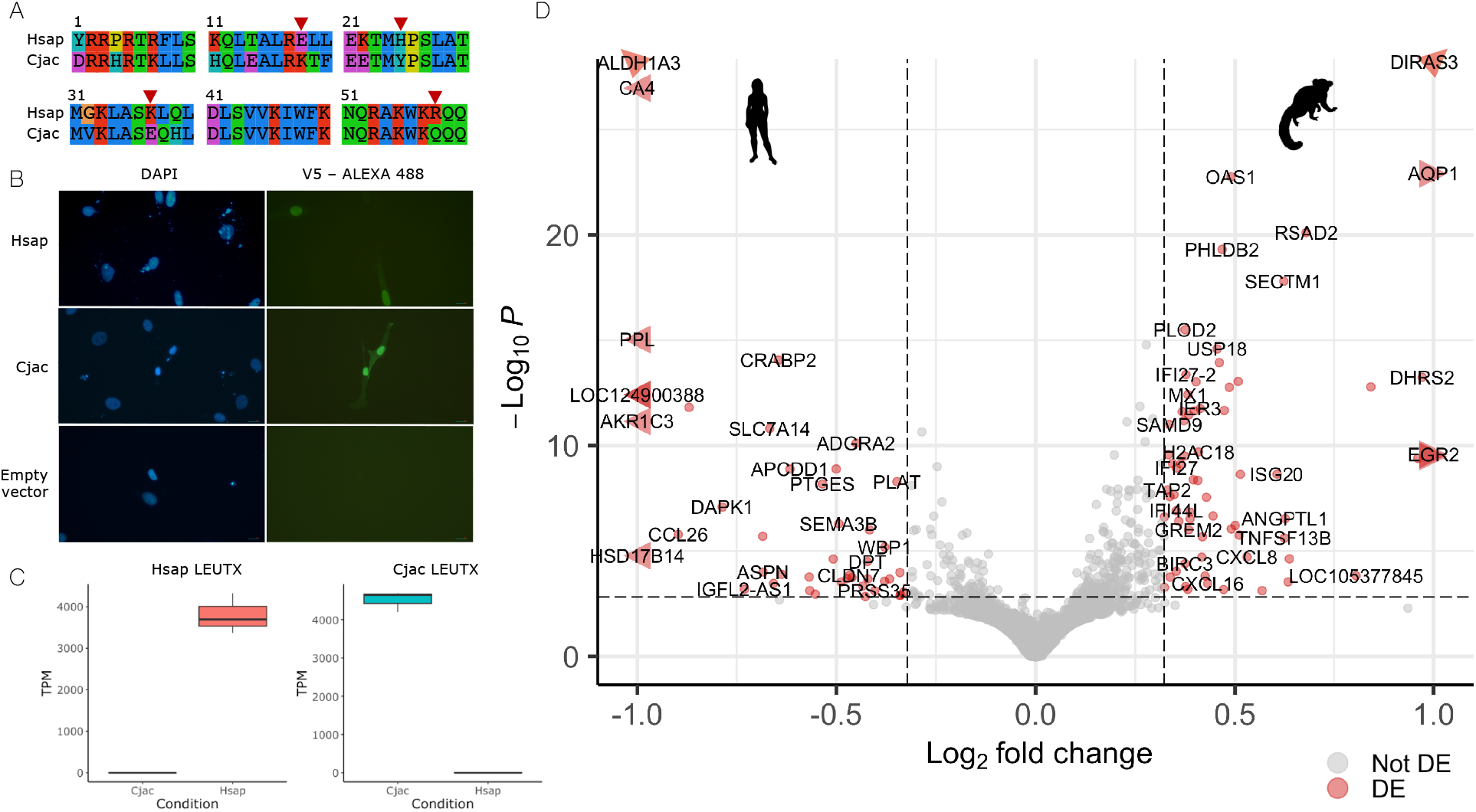
Ectopic expression of human and marmoset *LEUTX* genes. **A)** Alignment of LEUTX HD sequences of the species used in ectopic expression experiments. Red arrowheads mark residues under positive selection with a substitution between human and marmoset. **B)** Immunocytochemistry of fibroblasts transfected with human and marmoset *LEUTX* genes. DAPI stains DNA blue in cell nuclei. Expression of V5-tagged LEUTX proteins is detected with anti-V5 primary antibodies and Alexa Fluor 488 (green fluorescence)-labelled secondary antibodies. Empty vector transfections showed no green fluorescence. **C)** Expression of transfected *LEUTX* genes in cell culture samples. **D)** Transcriptional response to expression of human and marmoset *LEUTX* genes. Points to the left of centre are genes more highly expressed in response to human *LEUTX* expression than marmoset; points to the right are genes more highly expressed in response to marmoset *LEUTX* expression. DE genes (adjusted *P* < 0.05 and fold change > 1.25) are labelled with gene IDs and shown in red. Abbreviations: Cjac = *Callithrix jacchus*; DE = differentially expressed; Hsap = *Homo sapiens*; TPM = transcripts per million.

RNA-sequencing was performed on three biological replicates for human and marmoset *LEUTX*, and gene expression was then quantified with Kallisto (Bray et al. 2016) (Supplementary Table S3). Human (mean transcripts per million [TPM] = 3797) and marmoset (mean TPM = 4514) *LEUTX* genes were successfully expressed in the expected samples (Figure 5C). Differential expression analysis was performed to identify differences in the downstream genes responding to human versus marmoset *LEUTX*. We found that expression of human and marmoset *LEUTX* caused small but notable differences in the transcriptomic response within the transfected cells: 68 genes were more highly expressed in the marmoset-transfected samples, and 44 more highly expressed in the human-transfected samples (Figure 5D; Supplementary Tables S4; S5). Previous work found expression of human LEUTX to downregulate 754 and upregulate 481 genes (Maeso et al. 2016); this suggests that approximately 9% of the transcriptomic response to human LEUTX is different when marmoset LEUTX is expressed.

We sought to understand the significance of these transcriptional differences. We find that of the 68 genes more highly expressed in the marmoset treatment compared to human treatment, 33 were previously shown to be downregulated by human *LEUTX* (Maeso et al. 2016) (Figure 6A). This suggests that some genes downregulated by human LEUTX are not downregulated (or significantly less so) by marmoset LEUTX, revealing a change in transcription factor function. We performed biological process gene ontology (GO) analysis on these 68 DE genes: all of the top 20 GO terms without exception relate to the response to external biotic stimuli (Figure 6B; Supplementary Table S6). These terms do not appear in the gene set more highly expressed in response to human *LEUTX* than marmoset (Supplementary Table S7).

**Fig. 6.**
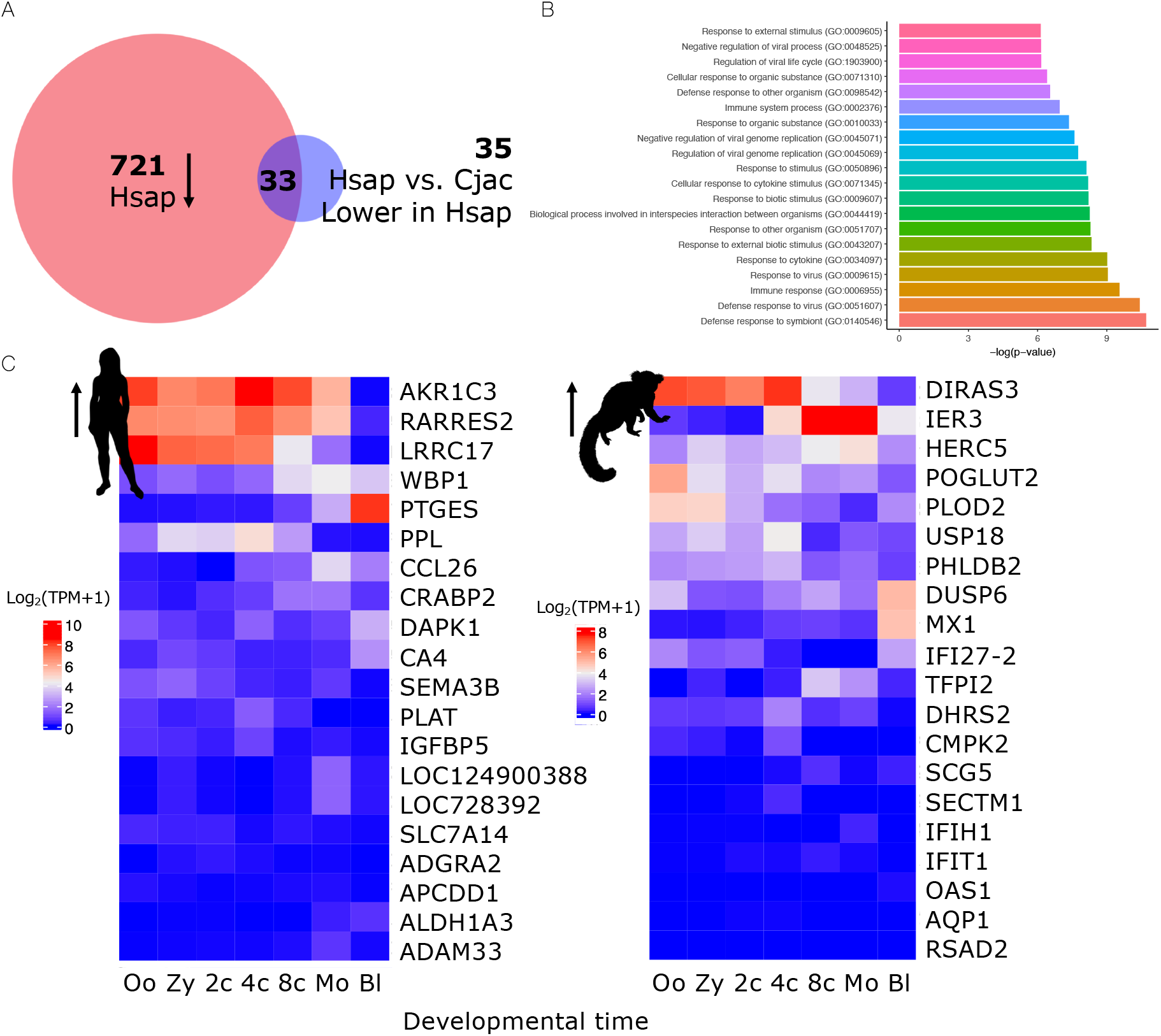
Differing transcriptional responses to human and marmoset LEUTX expression. **A)** Of the 68 genes more highly expressed in the marmoset LEUTX treatment compared to human LEUTX, 33 are known to be downregulated by human LEUTX. This suggests that they are not (or significantly less) downregulated by marmoset LEUTX. **B)** Top 20 GO terms enriched in the gene set upregulated in response to marmoset LEUTX compared to human LEUTX. **C)** Expression of DE genes in the human preimplantation embryo. Left heatmap shows top 20 genes upregulated in response to human LEUTX compared to marmoset, right heatmap shows top 20 genes upregulated in response to marmoset LEUTX compared to human. Abbreviations: 2c = two-cell; 4c = four-cell; 8c = 8-cell; Bl = blastocyst; Cjac = *Callithrix jacchus*; Hsap = *Homo sapiens*; Mo = morula; Oo = oocyte; TPM = transcripts per million; Zy = zygote.

We also explored the genes with the strongest difference in response to ectopic expression of human and marmoset *LEUTX*. This allowed us to test whether the genes identified by the ectopic expression approach are realistic embryonic targets of *LEUTX*. We profiled the temporal expression of the top 20 DE genes using published transcriptomic data (Yan et al. 2013) spanning human preimplantation development (Figure 6C; Supplementary Table S8). Ten of the top 20 genes upregulated in the human treatment compared to marmoset are expressed (TPM > 2 in at least one embryonic stage) during preimplantation development. Similarly, 12 of the 20 genes most differentially upregulated in response to marmoset LEUTX are embryonic genes. In addition, several of the most highly expressed DE genes (*AKR1C3*, *RARRES2*, *LRRC17*, *DIRAS3*) are strongly downregulated around the 8-cell to early morula stage, consistent with the timing of *LEUTX* expression. This suggests that the downstream targets differentially affected by marmoset and human *LEUTX* are realistic in vivo targets of this homeodomain protein.

Overall, there are clear, significant but relatively minor differences in the downstream targets of human and marmoset LEUTX, suggesting that the evolution of *LEUTX* sequences within the primate lineage has served to subtly modify the proteins’ transcription factor function rather than elicit dramatic shifts in target gene sets.

## Discussion

Fast-evolving homeobox genes may have received less attention than their highly conserved counterparts, but it is becoming increasingly clear that they play important roles in early embryonic development in mammals (Holland et al. 2017; MacLean & Wilkinson 2010; Niu et al. 2011; Maeso et al. 2016; Madissoon et al. 2016). One group with known roles in preimplantation development is ETCHbox, a set of genes in which copy number, protein-coding sequence, and protein functions have been shown to vary greatly between mammalian taxa (Maeso et al. 2016; Lewin et al. 2021; Royall et al. 2018). In this work, we characterised one of the ETCHbox genes within one taxonomic order, the primates, examining copy number, amino acid sequence evolution and divergence of protein function.

Comparative genomic analysis revealed that rapid evolution of the LEUTX protein-coding sequence has occurred to a remarkable extent within the primate lineage. While the CRX HD has remained almost completely unchanged, LEUTX has undergone divergence between primate clades, resulting in an amino acid sequence identity of only 35% between the two most divergent LEUTX HDs in our study, and an average of 70% across all sampled HDs. Positive selection acting on *LEUTX* sequences played an important role in this divergence, acting on key residues across the protein, including six within the HD. The most notable target of positive selection is residue 58, known to be a factor in determining the specificity of LEUTX proteins (Katayama et al. 2018), suggesting there has been selection for changes in protein targets.

Structural modelling revealed that targets of positive selection at HD positions 18 and 37 are positioned on the exterior of their respective helices. A network of salt bridges is known to form between the surfaces of helices 1 and 2 to stabilise HD structure (Clarke et al. 1994). Salt bridges are bonds between oppositely charged glutamic acid or aspartic acid (negatively charged) and arginine or lysine (positively charged) residues which contribute to protein structure, stability and specificity (Donald et al. 2011; Bosshard et al. 2004). In the human LEUTX protein, the residues at positions 18 and 37 are glutamic acid and lysine, respectively; this suggests selection for modifications to salt bridge formation has occurred within primates. Previous work sampling more divergent mammals also found positive selection at these residues (Lewin et al. 2021), suggesting that they have been consistent targets for selection across the Eutheria.

*LEUTX* is not lost or pseudogenised in any of the sampled primate species, implying selection for its retention. Although a small number of duplications are observed, these are almost entirely limited to the Strepsirrhini. The relative stability of *LEUTX* copy number within primates is a notable contrast to the situation across mammals more widely, in which this gene has been lost on at least four independent occasions and has duplicated in multiple species (Lewin et al. 2021). It is enlightening to compare the scenario of *LEUTX* with that of the Reproductive homeobox (Rhox) and Double homeobox (Dux) gene families. Both Rhox and Dux families are PRD-like genes which are mammal-specific, expressed during early development and have rapidly evolving sequences (Leidenroth & Hewitt 2010; Eidahl et al. 2016; MacLean & Wilkinson 2010; MacLean et al. 2005). Like *LEUTX*, the *RHOXF2* protein-coding sequence has diverged rapidly between primates, and copy number variation facilitated by nearby endogenous retroviral sequences also exists between closely related species, such as the presence of two copies in human and six in chimpanzee (Niu et al. 2011). From Dux genes, we learn that the presence of rapid evolutionary change does not indicate a lack of functional importance, as mouse Dux and its human orthologue DUX4 are both central to EGA despite minimal sequence conservation (Macfarlan et al. 2012; Peaston et al. 2004; De Iaco et al. 2017; Hendrickson et al. 2017; Eidahl et al. 2016; Vuoristo et al. 2022; Yoshihara et al. 2022). The parallels between these three gene families support the idea that selection pressures are acting to drive the evolutionary divergence of groups of homeobox genes with key roles in preimplantation development.

Bioinformatic analyses can reveal evolutionary constraint and the action of positive selection, but do not alone reveal the functional consequences of these changes. Using ectopic expression in primary cells, we compared the downstream actions of human *LEUTX* to the orthologous gene in the common marmoset *Callithrix jacchus*. Differential expression analysis revealed that expression of human and marmoset LEUTX proteins elicits small but notable differences in transcriptomic response. While this stands in stark contrast to the striking differences observed in the function of ARGFX when it was compared across a larger phylogenetic distance between human and cattle (Lewin et al. 2022), it suggests that positive selection has driven minor but detectable changes in LEUTX target specificity between primate species.

What explains this divergence of protein function? LEUTX is a transcription factor activated at EGA with expression at a critical point of mammalian embryonic development (Jouhilahti et al. 2016). At a molecular level, the gene regulatory networks (GRNs) underlying early preimplantation development at the time of, and immediately following, *LEUTX* expression are largely similar across primates but do exhibit small differences (Xinyi Wang et al. 2017; Hu et al. 2021; Nakamura et al. 2016). For instance, expression of factors forming the core pluripotency network of the epiblast (NANOG, POU5F1 and SOX2) is conserved between human and marmoset, but further epiblast-specific factors such as CREB3L1, HEY2, INSR and VENTX are species-specific (Boroviak et al. 2018). Overall, the relatively minor differences in LEUTX function between human and marmoset are consistent with the small-scale divergence of the GRNs coordinating preimplantation development; this suggests that positive selection on LEUTX proteins is fine-tuning their roles, changing targets at the periphery of largely conserved GRNs rather than initiating whole-scale changes to the core factors. The observed small differences in timing of LEUTX expression, which is highly specific to the 8-cell stage in humans but expressed in both 4-cell and 8-cell blastomeres in the marmoset, also supports the conclusion that rapid sequence evolution has driven small functional adjustments within the primate order. However, such adjustments should not be disregarded as superficial; early development in human and marmoset do indeed entail notable differences, including the duration of preimplantation development and the manner of implantation (Siriwardena & Boroviak 2022; Carter & Enders 2004; Boroviak et al. 2018).

## Conclusions

*LEUTX* is a fast-evolving homeobox gene recruited to a role in EGA in the early mammalian embryo. We characterised the *LEUTX* loci of all available chromosome-level primate genome assemblies, revealing dramatic divergence of protein-coding sequences but limited copy number variation. This divergence has been driven at least in part by positive selection, and six residues in the LEUTX HD were identified as targets of selection within the primate lineage. Ectopic expression experiments suggest that evolutionary sequence change has led to a small divergence in *LEUTX* function between primate species.

## Materials and Methods

### Comparative genomics

All reference assemblies of primates with a scaffold N50 of at least 1 Mb were downloaded from NCBI Genome (www.ncbi.nlm.nih.gov/genome/), with selected other species added to improve taxon representation (Supplementary Table S1). *LEUTX* genes were identified using blastn and tblastn searches and synteny; gene trees and reciprocal blast searches were used to confirm gene identities. The full human *LEUTX* sequence determined from transcriptome data (Maeso et al. 2016) was used as the basis for inferring gene structures. Genes with a complete HD are considered putatively functional. Intronless genes (putative retrocopies) are likely to be non-functional due to the absence of regulatory elements (Hurles 2004) and are therefore excluded. In two species, *O. garnettii* and *N. bengalensis*, we were unable to identify the first exon of *LEUTX*.

For phylogenetics, the maximum likelihood (ML) algorithm of IQ-TREE (Nguyen et al. 2015) was run with 1000 bootstraps made using UFBoot2 (Hoang et al. 2018) and automated model selection by ModelFinder (Kalyaanamoorthy et al. 2017). Sequence alignments were made using Clustal Omega (Sievers et al. 2011) implemented in Seaview version 4.7 (Gouy et al. 2010). A species tree was made using TimeTree 5, which uses a global time-calibrated tree of life synthesised from 4,075 studies (Kumar et al. 2017, 2022). Homeodomain sequences of PRD-class proteins were obtained from HomeoDB (Zhong & Holland 2011; Zhong et al. 2008).

Branch-Site Unrestricted Statistical Test for Episodic Diversification (BUSTED) (Murrell et al. 2015) was used to test whether positive selection has acted on *LEUTX* within the primates. The mixed effects branch-site model of MEME (Murrell et al. 2012) was then used to infer sites at which positive selection has acted, and the Fixed Effects Likelihood (FEL) model used to identify pervasive purifying selection (residues where purifying selection is detectable across the whole tree) (Kosakovsky Pond & Frost 2005). Tests for selection were run with default parameters using Datamonkey (Weaver et al. 2018). Where species have a *LEUTX* duplication, only one gene was used in the tests. *Cercopithecus mona* and *Chlorocebus sabaeus* sequences were included up to the ancestral start codon even though this has been lost; their complete HD suggests them to be functional.

The protein structure of the LEUTX homeodomain was modelled by comparative structural modelling using UCSF Chimera 1.16 (Pettersen et al. 2004) to implement Modeller (Šali & Blundell 1993). The *Drosophila melanogaster* Aristaless (Al) homeodomain (PRD-class) in complex with DNA (RCSB Protein Data Bank entry 3LNQ) (Berman et al. 2000; Miyazono et al. 2010) was taken as a reference. Putative ubiquitination sites were detected with ESA-UbiSite (Jyun-Rong Wang et al. 2017). Homeodomain residues were excluded as potential sites of ubiquitination.

For expression analysis, raw RNA-seq reads from human (*Homo sapiens*; PRJNA153427) (Yan et al. 2013), rhesus macaque (*Macaca mulatta*; PRJNA401876) (Chitwood et al. 2017) and common marmoset (*Callithrix jacchus*; PRJEB29285) preimplantation embryos were obtained from NCBI BioProject (www.ncbi.nlm.nih.gov/bioproject/). *LEUTX* expression was quantified at each developmental stage using Kallisto version 0.48.0 (Bray et al. 2016). Heatmaps were made using ComplexHeatmap version 2.8.0 (Gu et al. 2016) in R version 4.1.0 (R Core Team 2021).

### Ectopic expression

Primary human dermal fibroblasts (HDFs) (Stemnovate #SV-HF21-17-500) were maintained in HDF medium at 37°C with 5% CO_2_ and passaged at approximately 70% confluency every 3-4 days. HDF medium consists of Dulbecco’s modified Eagle medium (Gibco #41965039) with 10% heat-inactivated fetal bovine serum (Gibco #10500064) and 1% penicillin-streptomycin (Gibco #15140122). Testing for mycoplasma (Sigma Aldrich #MP0035) revealed no contamination.

Codon-optimised sequences of *Homo sapiens* and *Callithrix jacchus LEUTX* with a GGGGSGGGGS linker and C-terminal V5 tag (Supplementary Figure S5) were synthesised by ThermoFisher GeneArt and cloned into a pcDNA3.1 mammalian expression vector. For transfection, 65000 cells per well were seeded into 6-well plates. After 16 hours (h), medium was replaced with 2 mL antibiotic-free HDF medium. For each biological replicate, 108 μL Opti-MEM (Gibco #31985-062) was combined with 9.6 μL FUGENE 6 (Promega #E2691) and incubated for 5 minutes, then 2.4 μL of 1 μg/μL appropriate expression construct added before another 15-minute incubation. To each well, 120 μL of this mixture was added, and cells kept at 37°C with 5% CO_2_. After 24 h, transfection medium was removed and replaced with 2 mL HDF medium with 800 μg/mL G418 selective antibiotic (Gibco #10131-035). At 48 h post-transfection, RNA was extracted using an RNeasy Plus Micro kit (Qiagen #74034), and integrity tested using an Agilent 2100 Bioanalyzer.

To confirm expression of full-length proteins, the immunocytochemistry protocol of Maeso et al. (2016) was used with minor modifications: primary antibody (V5 tag monoclonal antibody; Invitrogen #37-7500) 1:500, 4 h incubation; secondary antibody (goat anti-mouse IgG H + L superclonal recombinant secondary antibody with Alexa Fluor 488; Invitrogen #A28175) 1:1000, 1 h incubation. Cells were incubated with DAPI (Invitrogen #S36938) to label nuclei. Results were visualised with an Olympus CKX53 inverted fluorescence microscope.

### Analysis of RNA-seq data

Three replicates for each treatment were sequenced on the Illumina NovaSeq 6000 platform (Novogene). FastQC version 0.11.8 (Andrews 2010) and MultiQC version 1.8 (Ewels et al. 2016) were used for quality control, and reads (150 bp paired-end) were subjected to filtering to remove adapter-containing reads, low-quality reads (Qscore < 5), and reads with > 10% Ns (undetermined bases), resulting in an average of 45.8 million reads per sample. Pseudoalignment to the human transcriptome from genome build GRCh38.p14 (RefSeq annotation) was performed with Kallisto version 0.48.0 (Bray et al. 2016); pseudoalignments were found for an average of 93.8% of reads. Gene-level transcript abundance estimates were created using tximport version 1.20.0 (Soneson et al. 2016), and then differential expression analysis completed in DESeq2 version 1.32.0 (Love et al. 2014) using apeglm (Zhu et al. 2019) for log fold change (LFC) shrinkage. EnhancedVolcano (Blighe et al. 2022) version 1.16.0 was used to create volcano plots. Genes with an adjusted *p*-value < 0.05, fold change > 1.25 and mean TPM over 2 were considered differentially expressed. To check whether differentially expressed genes represented realistic embryonic targets, raw reads from human preimplantation development (PRJNA153427) (Yan et al. 2013) were quantified with Kallisto as above (Bray et al. 2016). Gene ontology (GO) analysis was performed using PANTHER version 17.0 (Thomas et al. 2022) with Fisher’s exact test and a false discovery rate (FDR) correction of 0.05.

## Acknowledgements

We would like to thank Peter Mulhair for productive discussions, advice and critical reading of the manuscript. This work was supported by funding from the Biotechnology and Biological Sciences Research Council (BBSRC) (grant number BB/M011224/1); an Oxford-Wolfson Marriott BBSRC Graduate Scholarship; and Merton College, Oxford.

## Data availability

Raw and processed sequencing datasets will be available from the NCBI Gene Expression Omnibus (www.ncbi.nlm.nih.gov/geo) (Accession TBC).

## References

Andrews S. 2010. FastQC: A Quality Control Tool for High Throughput Sequence Data. Babraham Institute. Available online at: https://www.bioinformatics.babraham.ac.uk/projects/fastqc/.

Berman HM et al. 2000. The Protein Data Bank. Nucleic Acids Res. 28:235–242. doi: 10.1093/nar/28.1.235.

Blighe K, Rana S, Lewis M. 2022. EnhancedVolcano: Publication-ready volcano plots with enhanced colouring and labelling. R package version 1.16.0. https://github.com/kevinblighe/EnhancedVolcano

Boroviak T et al. 2018. Single cell transcriptome analysis of human, marmoset and mouse embryos reveals common and divergent features of preimplantation development. Development. 45:dev167833. doi: 10.1242/dev.167833.

Bosshard HR, Marti DN, Jelesarov I. 2004. Protein stabilization by salt bridges: concepts, experimental approaches and clarification of some misunderstandings. J Mol Recognit. 17:1–16. doi: https://doi.org/10.1002/jmr.657.

Bray NL, Pimentel H, Melsted P, Pachter L. 2016. Near-optimal probabilistic RNA-seq quantification. Nat Biotechnol. 34:525–527. doi: 10.1038/nbt.3519.

Bürglin TR, Affolter M. 2016. Homeodomain proteins: an update. Chromosoma. 125:497–521. doi: 10.1007/s00412-015-0543-8.

Carroll SB. 2000. Endless Forms: The Evolution of Gene Regulation and Morphological Diversity. Cell. 101:577–580. doi: 10.1016/S0092-8674(00)80868-5.

Carroll SB. 2008. Evo-Devo and an Expanding Evolutionary Synthesis: A Genetic Theory of Morphological Evolution. Cell. 134:25–36. doi: 10.1016/j.cell.2008.06.030.

Carter AM, Enders AC. 2004. Comparative aspects of trophoblast development and placentation. Reprod Biol Endocrinol. 2:46. doi: 10.1186/1477-7827-2-46.

Chitwood JL, Burruel VR, Halstead MM, Meyers SA, Ross PJ. 2017. Transcriptome profiling of individual rhesus macaque oocytes and preimplantation embryos. Biol Reprod. 97:353–364. doi: 10.1093/biolre/iox114.

Clarke ND, Kissinger CR, Desjarlais J, Gilliland GL, Pabo CO. 1994. Structural studies of the engrailed homeodomain. Protein Sci. 3:1779–1787. doi: 10.1002/pro.5560031018.

Cui W et al. 2016. Towards Functional Annotation of the Preimplantation Transcriptome: An RNAi Screen in Mammalian Embryos. Sci Rep. 6:37396. doi: 10.1038/srep37396.

Donald JE, Kulp DW, DeGrado WF. 2011. Salt bridges: geometrically specific, designable interactions. Proteins. 79:898–915. doi: 10.1002/prot.22927.

Duboule D. 1994. Guidebook to the Homeobox Genes. Oxford: Oxford University Press.

Eidahl JO et al. 2016. Mouse Dux is myotoxic and shares partial functional homology with its human paralog DUX4. Hum Mol Genet. 25:4577–4589. doi: 10.1093/hmg/ddw287.

Ewels P, Magnusson M, Lundin S, Käller M. 2016. MultiQC: Summarize analysis results for multiple tools and samples in a single report. Bioinformatics. 32:3047–3048. doi: 10.1093/bioinformatics/btw354.

Gehring WJ, Affolter M, Bürglin T. 1994. Homeodomain proteins. Annu Rev Biochem. 63:487–526. doi: 10.1146/annurev.bi.63.070194.002415.

Gouy M, Guindon S, Gascuel O. 2010. SeaView version 4: A multiplatform graphical user interface for sequence alignment and phylogenetic tree building. Mol Biol Evol. 27:221–224. doi: 10.1093/molbev/msp259.

Gu Z, Eils R, Schlesner M. 2016. Complex heatmaps reveal patterns and correlations in multidimensional genomic data. Bioinformatics. 32:2847–2849. doi: 10.1093/bioinformatics/btw313.

Guo Y et al. 2022. Obox4 secures zygotic genome activation upon loss of Dux. bioRxiv. 2022.07.04.498763. doi: 10.1101/2022.07.04.498763.

Hendrickson PG et al. 2017. Conserved roles of mouse DUX and human DUX4 in activating cleavage-stage genes and MERVL/HERVL retrotransposons. Nat Genet. 49:925–934. doi: 10.1038/ng.3844.

Hoang DT, Chernomor O, von Haeseler A, Minh BQ, Vinh LS. 2018. UFBoot2: Improving the Ultrafast Bootstrap Approximation. Mol Biol Evol. 35:518–522. doi: 10.1093/molbev/msx281.

Holland PWH, Marlétaz F, Maeso I, Dunwell TL, Paps J. 2017. New genes from old: Asymmetric divergence of gene duplicates and the evolution of development. Philos Trans R Soc Lond B Biol Sci. 372:20150480. doi: 10.1098/rstb.2015.0480.

Hu Y et al. 2021. Single-cell analysis of nonhuman primate preimplantation development in comparison to humans and mice. Dev Dyn. 250:974–985. doi: 10.1002/dvdy.295.

Hurles M. 2004. Gene duplication: the genomic trade in spare parts. PLoS Biol. 2:e206.

De Iaco A et al. 2017. DUX-family transcription factors regulate zygotic genome activation in placental mammals. Nat. Genet. doi: 10.1038/ng.3858.

Jouhilahti E-M et al. 2016. The human PRD-like homeobox gene LEUTX has a central role in embryo genome activation. Development. 143:3459–3469. doi: 10.1242/dev.134510.

Kalyaanamoorthy S, Minh BQ, Wong TKF, von Haeseler A, Jermiin LS. 2017. ModelFinder: fast model selection for accurate phylogenetic estimates. Nat Methods. 14:587–589. doi: 10.1038/nmeth.4285.

Katayama S et al. 2018. Phylogenetic and mutational analyses of human LEUTX, a homeobox gene implicated in embryogenesis. Sci Rep. 8:17421. doi: 10.1038/s41598-018-35547-5.

Kosakovsky Pond SL, Frost SDW. 2005. Not So Different After All: A Comparison of Methods for Detecting Amino Acid Sites Under Selection. Mol Biol Evol. 22:1208–1222. doi: 10.1093/molbev/msi105.

Kumar S et al. 2022. TimeTree 5: An Expanded Resource for Species Divergence Times. Mol. Biol. Evol. 39:msac174. doi: 10.1093/molbev/msac174.

Kumar S, Stecher G, Suleski M, Hedges SB. 2017. TimeTree: A Resource for Timelines, Timetrees, and Divergence Times. Mol Biol Evol. 34:1812–1819. doi: 10.1093/molbev/msx116.

Leidenroth A, Hewitt JE. 2010. A family history of DUX4: phylogenetic analysis of DUXA, B, C and Duxbl reveals the ancestral DUX gene. BMC Evol Biol. 10:364. doi: 10.1186/1471-2148-10-364.

Lewin TD, Fouladi-Nashta AA, Holland PWH. 2022. PRD-Class Homeobox Genes in Bovine Early Embryos: Function, Evolution, and Overlapping Roles. Mol Biol Evol. 39:msac098. doi: 10.1093/molbev/msac098.

Lewin TD, Royall AH, Holland PWH. 2021. Dynamic Molecular Evolution of Mammalian Homeobox Genes: Duplication, Loss, Divergence and Gene Conversion Sculpt PRD-Class Repertoires. J Mol Evol. 89:396–414. doi: 10.1007/s00239-021-10012-6.

Li H et al. 2006. A novel maternally transcribed homeobox gene, Eso-1, is preferentially expressed in oocytes and regulated by cytoplasmic polyadenylation. Mol Reprod Dev. 73:825–833. doi: 10.1002/mrd.20478.

Love MI, Huber W, Anders S. 2014. Moderated estimation of fold change and dispersion for RNA-seq data with DESeq2. Genome Biol. 15:550. doi: 10.1186/s13059-014-0550-8.

Macfarlan TS et al. 2012. Embryonic stem cell potency fluctuates with endogenous retrovirus activity. Nature. 487:57–63. doi: 10.1038/nature11244.

MacLean JA et al. 2005. Rhox: A new homeobox gene cluster. Cell. 120:369–382. doi: 10.1016/j.cell.2004.12.022.

MacLean JA, Wilkinson MF. 2010. The Rhox genes. Reproduction. 140:195–213. doi: 10.1530/REP-10-0100.

Madissoon E et al. 2016. Characterization and target genes of nine human PRD-like homeobox domain genes expressed exclusively in early embryos. Sci Rep. 6:28995. doi: 10.1038/srep28995.

Maeso I et al. 2016. Evolutionary origin and functional divergence of totipotent cell homeobox genes in eutherian mammals. 14:45. BMC Biol. doi: 10.1186/s12915-016-0267-0.

Mazid MA et al. 2022. Rolling back of human pluripotent stem cells to an 8-cell embryo-like stage. Nature. 605:315–324. doi: 10.1038/s41586-022-04625-0.

Miyazono K-I et al. 2010. Cooperative DNA-binding and sequence-recognition mechanism of aristaless and clawless. EMBO J. 29:1613–23. doi: 10.1038/emboj.2010.53.

Murrell B et al. 2012. Detecting individual sites subject to episodic diversifying selection. PLoS Genet. 8:e1002764. doi: 10.1371/journal.pgen.1002764.

Murrell B et al. 2015. Gene-wide identification of episodic selection. Mol Biol Evol. 32:1365–1371. doi: 10.1093/molbev/msv035.

Nakamura T et al. 2016. A developmental coordinate of pluripotency among mice, monkeys and humans. Nature. 537:57–62. doi: 10.1038/nature19096.

Nguyen L-T, Schmidt HA, von Haeseler A, Minh BQ. 2015. IQ-TREE: A Fast and Effective Stochastic Algorithm for Estimating Maximum-Likelihood Phylogenies. Mol Biol Evol. 32:268–274. doi: 10.1093/molbev/msu300.

Niu AL et al. 2011. Rapid evolution and copy number variation of primate RHOXF2, an X-linked homeobox gene involved in male reproduction and possibly brain function. BMC Evol Biology. 11:298. doi: 10.1186/1471-2148-11-298.

Peaston AE et al. 2004. Retrotransposons regulate host genes in mouse oocytes and preimplantation embryos. Dev Cell. 7:597–606. doi: 10.1016/j.devcel.2004.09.004.

Pettersen EF et al. 2004. UCSF Chimera - A visualization system for exploratory research and analysis. J Comput Chem. 25:1605–1612. doi: 10.1002/jcc.20084.

Piskacek S et al. 2007. Nine-amino-acid transactivation domain: Establishment and prediction utilities. Genomics. 89:756–768. doi: 10.1016/j.ygeno.2007.02.003.

Pozzi L et al. 2014. Primate phylogenetic relationships and divergence dates inferred from complete mitochondrial genomes. Mol Phylogenet Evol. 75:165–183. doi: 10.1016/j.ympev.2014.02.023.

R Core Team. 2021. R: A language and environment for statistical computing. R Foundation for Statistical Computing, Vienna, Austria. URL https://www.R-project.org/.

dos Reis M et al. 2018. Using Phylogenomic Data to Explore the Effects of Relaxed Clocks and Calibration Strategies on Divergence Time Estimation: Primates as a Test Case. Syst Biol. 67:594–615. doi: 10.1093/sysbio/syy001.

Royall AH, Maeso I, Dunwell TL, Holland PWH. 2018. Mouse Obox and Crxos modulate preimplantation transcriptional profiles revealing similarity between paralogous mouse and human homeobox genes. Evodevo. 9:2. doi: 10.1186/s13227-018-0091-4.

Šali A, Blundell TL. 1993. Comparative protein modelling by satisfaction of spatial restraints. J Mol Biol. 234:779–815. doi: 10.1006/jmbi.1993.1626.

Sievers F et al. 2011. Fast, scalable generation of high-quality protein multiple sequence alignments using Clustal Omega. Mol Syst Biol. 7:539. doi: 10.1038/msb.2011.75.

Siriwardena D, Boroviak TE. 2022. Evolutionary divergence of embryo implantation in primates. Philos Trans R Soc Lond B Biol Sci. 377:20210256. doi: 10.1098/rstb.2021.0256.

Soneson C, Love MI, Robinson MD. 2016. Differential analyses for RNA-seq: Transcript-level estimates improve gene-level inferences. F1000Res. 4:1521. doi: 10.12688/F1000RESEARCH.7563.2.

Thomas PD et al. 2022. PANTHER: Making genome-scale phylogenetics accessible to all. Protein Sci. 31:8–22. doi: 10.1002/pro.4218.

Töhönen V et al. 2015. Novel PRD-like homeodomain transcription factors and retrotransposon elements in early human development. Nat Commun. 6:8207. doi: 10.1038/ncomms9207.

Vuoristo S et al. 2022. DUX4 is a multifunctional factor priming human embryonic genome activation. iScience. 25:104137. doi: 10.1016/j.isci.2022.104137.

Wang Jyun-Rong et al. 2017. ESA-UbiSite: accurate prediction of human ubiquitination sites by identifying a set of effective negatives. Bioinformatics. 33:661–668. doi: 10.1093/bioinformatics/btw701.

Wang Xinyi et al. 2017. Transcriptome analyses of rhesus monkey preimplantation embryos reveal a reduced capacity for DNA double-strand break repair in primate oocytes and early embryos. Genome Res. 27:567–579. doi: 10.1101/gr.198044.115.

Weaver S et al. 2018. Datamonkey 2.0: A modern web application for characterizing selective and other evolutionary processes. Mol Biol Evol. 35:773–777. doi: 10.1093/molbev/msx335.

Wilkinson RD et al. 2011. Dating Primate Divergences through an Integrated Analysis of Palaeontological and Molecular Data. Syst Biol. 60:16–31. doi: 10.1093/sysbio/syq054.

Yan L et al. 2013. Single-cell RNA-Seq profiling of human preimplantation embryos and embryonic stem cells. Nat Struct Mol Biol. 20:1131–1139. doi: 10.1038/nsmb.2660.

Yoshihara M et al. 2022. Transient DUX4 expression in human embryonic stem cells induces blastomere-like expression program that is marked by SLC34A2. Stem Cell Reports. 17:1743–1756. doi: 10.1016/j.stemcr.2022.06.002.

Zhong YF, Butts T, Holland PWH. 2008. HomeoDB: A database of homeobox gene diversity. Evol. Dev. 10:516–518. doi: 10.1111/j.1525-142X.2008.00266.x.

Zhong YF, Holland PWH. 2011. HomeoDB2: Functional expansion of a comparative homeobox gene database for evolutionary developmental biology. Evol Dev. 13:567–568. doi: 10.1111/j.1525-142X.2011.00513.x.

Zhu A, Ibrahim JG, Love MI. 2019. Heavy-tailed prior distributions for sequence count data: removing the noise and preserving large differences. Bioinformatics. 35:2084–2092. doi: 10.1093/bioinformatics/bty895.

Zou Z et al. 2022. Translatome and transcriptome co-profiling reveals a role of TPRXs in human zygotic genome activation. Science. 378:abo7923. doi: 10.1126/science.abo7923.

