## Supplementary Figures S1 - S5 for "Rapid evolution of the embryonically-expressed homeobox gene *LEUTX* within primates"

**Electronic Supplementary Material – Supplementary Figures S1 – S5**

**Fig. S1**

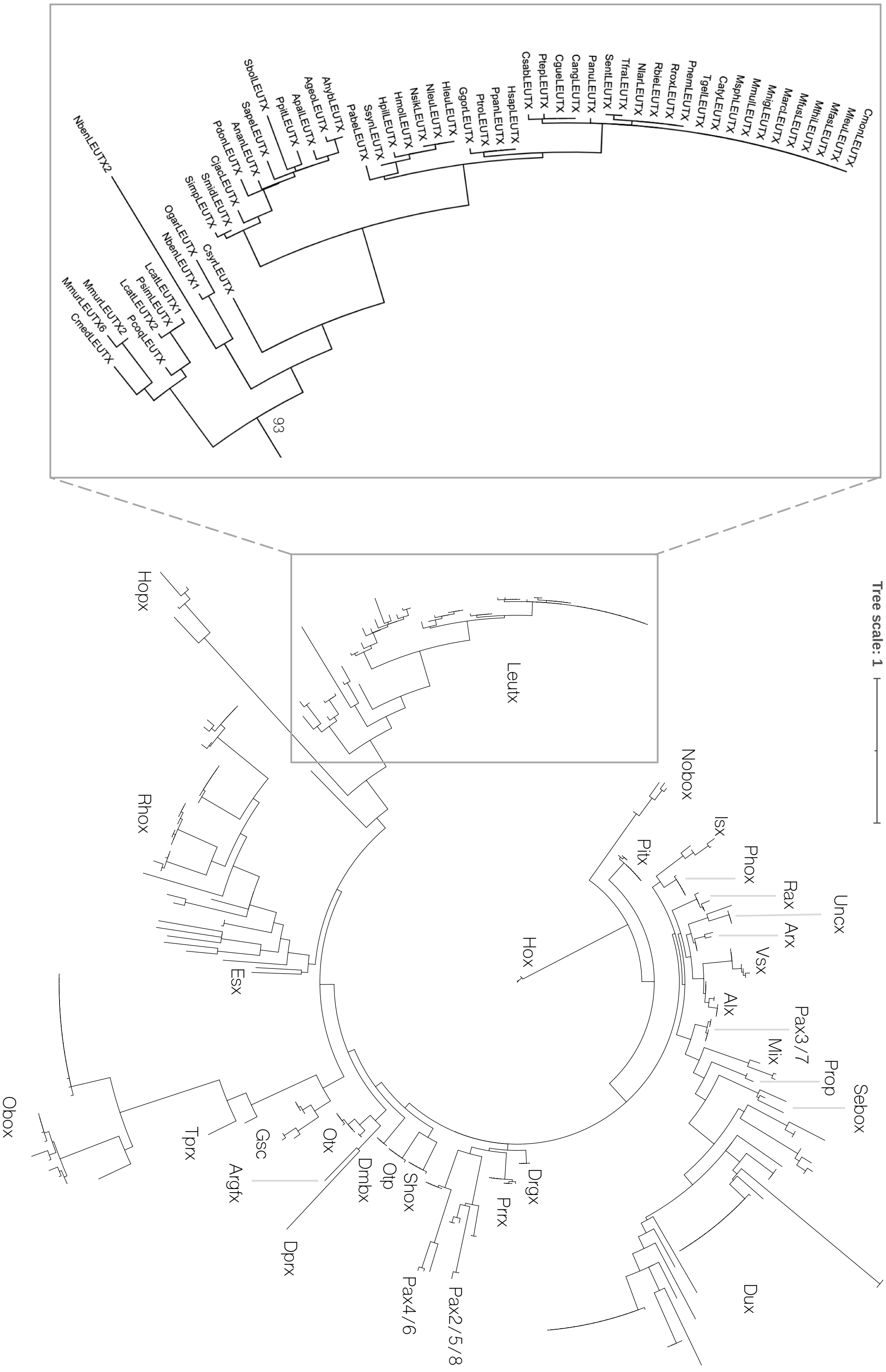
NbenLEUTX2
SbolLEUTX
SapeLEUTX
AnanLEUTX
PdonLEUTX
CiacLEUTX
SmidLEUTX
SimpLEUTX
PpilLEUTX
ApalLEUTX
AgeoLEUTX
AhybLEUTX
PabeLEUTX
SsynLEUTX
HpilLEUTX
HmolLEUTX
NsikLEUTX
NleuLEUTX
HleuLEUTX
GgorLEUTX
PtroLEUTX
PpanLEUTX
HsapLEUTX
CsabLEUTX
PtepLEUTX
CgueLEUTX
CangLEUTX
PanuLEUTX
SentLEUTX
TfraLEUTX
NlarLEUTX
RbieLEUTX
RroxLEUTX
PnemLEUTX
TgelLEUTX
CatyLEUTX
MsphLEUTX
MmulLEUTX
MnigLEUTX
MarcLEUTX
MfusLEUTX
MthiLEUTX
MfasLEUTX
MleuLEUTX
CmonLEUTX
OgarLEUTX
CsyrLEUTX
NbenLEUTX1
LcatLEUTX1
PsimLEUTX
LcatLEUTX2
PcoqLEUTX
MmurLEUTX2
MmurLEUTX6
CmedLEUTX
93
Hopx
Leutx
Tree scale: 1
Nobox
Rhox
Isx
Pitx
Phox
Rax
Uncx
Arx
Hox
Vsx
Esx
Alx
Pax3/7
Mix
Prop
Sebox
Obox
Tprx
Gsc
Otx
Argfx
Dmbx
Otp
Shox
Drgx
Prrx
Dprx
Dux
Pax4/6
Pax2/5/8

**Fig. S1 Position of LEUTX sequences within a tree of all PRD-class homeodomains.** Homeodomain sequences of PRD-class proteins were obtained from HomeoDB (Zhong et al. 2008; Zhong and Holland 2011). Species name abbreviations as in Fig. 1 main text.

### Fig. S2

```
            1
OgarCRX  MMAYMNPGPH YSVNALALSG PSVDLMHQPV PYPSAPRKQR RERTTFTRSQ LEELEALFAK TQYPDVYARE EVALKINLPE SRVQVWFKNR
PcoqCRX  MMAYMNPGPH YSVNALALSG PSVDLMHQAV PYPSAPRKQR RERTTFTRSQ LEELEALFAK TQYPDVYARE EVALKINLPE SRVQVWFKNR
MmurCRX  MMAYMNPGPH YSVNALALSG PSVDLVHQAV PYPSAPRKQR RERTTFTRSQ LEELEALFAK TQYPDVYARE EVALKINLPE SRVQVWFKNR
CsyrCRX  MMAYMNPGPH YSVNALALSG PSVDLMHPAV PYPSAPRKQR RERTTFTRSQ LEELEALFAK TQYPDVYARE EVALKINLPE SRVQVWFKNR
SapeCRX  MMAYMNPGPH YSVNTLALSG PSVDLMHQAV PYPSAPRKQR RERTTFTRSQ LEELEALFAK TQYPDVYARE EVALKINLPE SRVQVWFKNR
SbolCRX  MMAYMNPGPH YSVNTLALSG PSVDLMHQAV PYPSAPRKQR RERTTFTRSQ LEELEALFAK TQYPDVYARE EVALKINLPE SRVQVWFKNR
AnanCRX  MMAYMNPGPH YSVNTLALSG PSVDLMHQAV PYPSAPRKQR RERTTFTRSQ LEELEALFAK TQYPDVYARE EVALKINLPE SRVQVWFKNR
CimiCRX  MMAYMNPGPH YSVNTLALSG PSVDLMHQAV PYPSAPRKQR RERTTFTRSQ LEELEALFAK TQYPDVYARE EVALKINLPE SRVQVWFKNR
GgorCRX  MMAYMNPGPH YSVNALALSG PSVDLMHQAV PYPSAPRKQR RERTTFTRSQ LEELEALFAK TQYPDVYARE EVALKINLPE SRVQVWFKNR
HsapCRX  MMAYMNPGPH YSVNALALSG PSVDLMHQAV PYPSAPRKQR RERTTFTRSQ LEELEALFAK TQYPDVYARE EVALKINLPE SRVQVWFKNR
PtroCRX  MMAYMNPGPH YSVNALALSG PSVDLMHQAV PYPSAPRKQR RERTTFTRSQ LEELETLFAK TQYPDVYARE EVALKINLPE SRVQVWFKNR
CangCRX  MMAYMNPGPH YSVNALALSG PSVDLMHQAV PYPSAPRKQR RERTTFTRSQ LEELEALFAK TQYPDVYARE EVALKINLPE SRVQVWFKNR
PtepCRX  MMAYMNPGPH YSVNALALSG PSVDLMHQAV PYPSAPRKQR RERTTFTRSQ LEELEALFAK TQYPDVYARE EVALKINLPE SRVQVWFKNR
CsabCRX  MMAYMNPGPH YSVNALALSG PSVDLMHQAV PYPSAPRKQR RERTTFTRSQ LEELEALFAK TQYPDVYARE EVALKINLPE SRVQVWFKNR
TgelCRX  MMAYMNPGPH YSVNALALSG PSVDLMHQAV PYPSAPRKQR RERTTFTRSQ LEELEALFAK TQYPDVYARE EVALKINLPE SRVQVWFKNR
PanuCRX  MMAYMNPGPH YSVNALALSG PSVDLMHQAV PYPSAPRKQR RERTTFTRSQ LEELEALFAK TQYPDVYARE EVALKINLPE SRVQVWFKNR
CatyCRX  MMAYMNPGPH YSVNALALSG PSVDLMHQAV PYPSAPRKQR RERTTFTRSQ LEELEALFAK TQYPDVYARE EVALKINLPE SRVQVWFKNR
MleuCRX  MMAYMNPGPH YSVNALALSG PSVDLMHQAV PYPSAPRKQR RERTTFTRSQ LEELEALFAK TQYPDVYARE EVALKINLPE SRVQVWFKNR
MmulCRX  MMAYMNPGPH YSVNALALSG PSVDLMHQAV PYPSAPRKQR RERTTFTRSQ LEELEALFAK TQYPDVYARE EVALKINLPE SRVQVWFKNR
MnemCRX  MMAYMNPGPH YSVNALALSG PSVDLMHQAV PYPSAPRKQR RERTTFTRSQ LEELEALFAK TQYPDVYARE EVALKINLPE SRVQVWFKNR

        91
OgarCRX  RAKCRQQRQQ QKQQQQPPGG QAKARPAKRK AGTSPRLSTD VCPDPLGISD SYSPPLPGPS ASPTTAVATV SIWSPASESP LPEAQRAGLV
PcoqCRX  RAKCRQQRQQ QKQQQQPPGG QAKARPAKRK AGSSPRPSTD VCPDPLGISD SYSPPLPGPS GSPTTAVATV SIWSPASESP LPEAQRAGLV
MmurCRX  RAKCRQQRQQ QKQQQQPPGG QAKARPAKRK AGSSPRPSTD VCPDPLGISD SYSPPLPGPS GSPTTAVATV SIWSPASESP LPEAQRAGLV
CsyrCRX  RAKCRQQRQQ QKQQQQPPGG QAKARPAKRK AGTSPRPSTD VCPDPLGISD SYSPPLPGPS GSPTTAVATV SIWSPASESP LPEAQRAGLV
SapeCRX  RAKCRQQRQQ QKQQQQPPGS QAKARPAKRK AGTSPRPSTD VCPDPLGISD SYSPPLPGPS GSPSTAVATV SIWSPASESP LPEAQRAGLV
SbolCRX  RAKCRQQRQQ QKQQQQPPGG QAKARPAKRK AGTSPRPSTD VCPDPLGISD SYSPPLPGPS GSPSTAVATV SIWSPASESP LPEAQRAGLV
AnanCRX  RAKCRQQRQQ QKQQQQPPGG QAKARPAKRK AGTSPRPSTD VCPDPLGISD SYSPPLPGPS ASPNTAVATV SIWSPASESP LPEAQRAGLV
CimiCRX  RAKCRQQRQQ QKQQQQPPGS QAKARPAKRK AGTSPRPSTD VCPDPLGISD SYSPPLPGPS GSPSTAVATV SIWSPASESP LPEAQRAGLV
GgorCRX  RAKCRQQRQQ QKQQQQPPGG QAKARPAKRK AGTSPRPSTD VCPDPLGISD SYSPPLPGPS GSPTTAVATV SIWSPASESP LPEAQRAGLV
HsapCRX  RAKCRQQRQQ QKQQQQPPGG QAKARPAKRK AGTSPRPSTD VCPDPLGISD SYSPPLPGPS GSPTTAVATV SIWSPASESP LPEAQRAGLV
PtroCRX  RAKCRQQRQQ QKQQQQPPGG QAKARPAKRK AGTSPRPSTD VCSDPLGISD SYSPPLPGPS GSPTTAVATV SIWSPASESP LPEAQRAGLV
CangCRX  RAKCRQQRQQ QKQQQQPPGG QAKARPAKRK AGTSPRPSTD VCPDPLGISD SYSPPLPSPS GSPTTAVATV SIWSPASESP LPEVQRAGLV
PtepCRX  RAKCRQQRQQ QKQQQQPPGG QAKARPAKRK AGTSPRPSTD VCPDPLGISD SYSPPLPGPS GSPTTAVATV SIWSPASESP LPEAQRAGLV
CsabCRX  RAKCRQQRQQ QKQQQQPPGG QAKARPAKRK AGTSPRPSTD VCPDPLGISD SYSPPLPGPS GSPTTAVATV SIWSPASESP LPEAQRAGLV
TgelCRX  RAKCRQQRQQ QKQQQQPPGG QAKARPAKRK AGTSPRPSTD VCPDPLGISD SYSPPLPGPS GSPTTAVATV SIWSPASESP LPEAQRAGLV
PanuCRX  RAKCRQQRQQ QKQQQQPPGG QAKARPAKRK AGTSPRPSTD VCPDPLGISD SYSPPLPGPS GSPTTAVATV SIWSPASESP LPEAQRAGLV
CatyCRX  RAKCRQQRQQ QKQQQQPPGG QAKARPAKRK AGTSPRPSTD VCPDPLGISD SYSPPLPGPS GSPTTAVATV SIWSPASESP LPEAQRAGLV
MleuCRX  RAKCRQQRQQ QKQQQQPPGG QAKARPAKRK AGTSPRPSTD VCPDPLGISD SYSPPLPGPS GSPTTAVATV SIWSPASESP LPEAQRAGLV
MmulCRX  RAKCRQQRQQ QKQQQQPPGG QAKARPAKRK AGTSPRPSTD VCPDPLGISD SYSPPLPGPS GSPTTAVATV SIWSPASESP LPEAQRAGLV
MnemCRX  RAKCRQQRQQ QKQQQQPPGG QAKARPAKRK AGTSPRPSTD VCPDPLGISD SYSPPLPGPS GSPTTAVATV SIWSPASESP LPEAQRAGLV

       181
OgarCRX  ASGPSLTSAP YAMTYTPASA FCSSPSAYGS PSSYFSGLDP YLSPMVPQLG GPALSPLSGP SVGPSLAQSP TSLSGQSYGA YSPVDSLEFK
PcoqCRX  ASGPSLTSAP YAMTYAPASA FCSSPSAYGS PSSYFSGLDP YLSPMVPQLG GPALSPLSGP SVGPSLAQSP TSLSGQSYGP YSPVDSLEFK
MmurCRX  ASGPSLTSAP YAMTYAPASA FCSSPSAYGS PSSYFSGLDP YLSPMVPQLG GPALSPLSGP SVGPSLAQSP TSLSGQSYGP YSPVDSLEFK
CsyrCRX  ASGPSLTSAP YAMTYAPASA FCSSPSAYGS PSSYFSGLDP YLSPMVPQLG GPALSPLSGP SVGPSLAQSP TSLSGQSYSA YSPVDSLEFK
SapeCRX  ASAPSLTSAP YAMTYAPASA FCSSPSAYGS PSSYFSGLDP YLSPMVPQLG GPALSPLSGP SVGPSLAQSP TSLSGQSYGA YSPVDSLEFK
SbolCRX  ASAPSLTSAP YAMTYAPASA FCSSPSAYGS PSSYFSGLDP YLSPMVPQLG GPALSPLSGP SVGPSLAQSP TSLSGQSYGA YSPVDSLEFK
AnanCRX  ASAPSLTSAP YAMTYAPASA FCSSPSAYGS PSSYFSGLDP YLSPMVPQLG GPALSPLSGP SVGPSLAQSP TSLSGQSYGA YSPVDSLEFK
CimiCRX  ASAPSLTSAP YAMTYAPASA FCSSPSAYGS PSSYFSGLDP YLSPMVPQLG GPALSPLSGP SVGPSLAQSP TSLSGQSYGA YSPVDSLEFK
GgorCRX  ASGPSLTSAP YAMTYAPASA FCSSPSAYGS PSSYFSGLDP YLSPMVPQLG GPALSPLSGP SVGPSLAQSP TSLSGQSYGA YSPVDSLEFK
HsapCRX  ASGPSLTSAP YAMTYAPASA FCSSPSAYGS PSSYFSGLDP YLSPMVPQLG GPALSPLSGP SVGPSLAQSP TSLSGQSYGA YSPVDSLEFK
PtroCRX  ASGPSLTSAP YAMTYAPASA FCSSPSAYGS PSSYFSGLDP YLSPMVPQLG GPALSPLSGP SVGPSLAQSP TSLSGQSYGA YSPVDSLEFK
CangCRX  ASGPSLTSAP YAMTYAPASA FCSSPSAYGS PSSYFSSLDP YLSPMVPQLG GPALSPLSGP SVGPSLAQSP TSLSGQSYGA YSPVDSLEFK
PtepCRX  ASGPSLTSAP YAMTYAPASA FCSSPSAYGS PSSYFSGLDP YLSPMVPQLG GPALSPLSGP SVGPSLAQSP TSLSGQSYGA YSPVDSLEFK
CsabCRX  ASGPSLTSAP YAMTYAPASA FCSSPSAYGS PSSYFSGLDP YLSPMVPQLG GPALSPLSGP SVGPSLAQSP TSLSGQSYSA YSPVDSLEFK
TgelCRX  ASGPSLTSAP YAMTYAPASA FCSSPSAYGS PSSYFSGLDP YLSPMVPQLG GPALSPLSGP SVGPSLAQSP TSLSGQSYGA YSPVDSLEFK
PanuCRX  ASGPSLTSAP YAMTYAPASA FCSSPSAYGS PSSYFSGLDP YLSPMVPQLG GPALSPLSGP SVGPSLAQSP TSLSGQSYGA YSPVDSLEFK
CatyCRX  ASGPSLTSAP YAMTYAPASA FCSSPSAYGS PSSYFSGLDP YLSPMVPQLG GPALSPLSGP SVGPSLAQSP TSLSGQSYGA YSPVDSLEFK
MleuCRX  ASGPSLTSAP YAMTYAPASA FCSSPSAYGS PSSYFSGLDP YLSPMVPQLG GPALSPLSGP SVGPSLAQSP TSLSGQSYGA YSPVDSLEFK
MmulCRX  ASGPSLTSAP YAMTYAPASA FCSSPSAYGS PSSYFSGLDP YLSPMVPQLG GPALSPLSGP SVGPSLAQSP TSLSGQSYGA YSPVDSLEFK
MnemCRX  ASGPSLTSAP YAMTYAPASA FCSSPSAYGS PSSYFSGLDP YLSPMVPQLG GPALSPLSGP SVGPSLAQSP TSLSGQSYGA YSPVDSLEFK

       271
OgarCRX  DPSGTWKFTY NPMDPLDYKD QSAWKFQIL
PcoqCRX  DPSGTWKFTY NPMDPLDYKD QSAWKFQIL
MmurCRX  DPSGTWKFTY NPMDPLDYKD QSAWKFQIL
CsyrCRX  DPTGTWKFTY NPMDPLDYKD QSAWKFQIL
SapeCRX  DPTGTWKFTY NPMDPLDYKD QSAWKFQIL
SbolCRX  DPTGTWKFTY NPMDPLDYKD QSAWKFQIL
AnanCRX  DPTGTWKFTY NPMDPLDYKD QSAWKFQIL
CimiCRX  DPTGTWKFTY NPMDPLDYKD QSAWKFQIL
GgorCRX  DPTGTWKFTY NPMDPLDYKD QSAWKFQIL
HsapCRX  DPTGTWKFTY NPMDPLDYKD QSAWKFQIL
PtroCRX  DPTGTWKFTY NPMDPLDYKD QSAWKFQIL
CangCRX  DPTGTWKFTY NPMDPLDYKD QSAWKFQIL
PtepCRX  DPTGTWKFTY NPMDPLDYKD QSAWKFQIL
CsabCRX  DPTGTWKFTY NPMDPLDYKD QSAWKFQIL
TgelCRX  DPTGTWKFTY NPMDPLDYKD QSAWKFQIL
PanuCRX  DPTGTWKFTY NPMDPLDYKD QSAWKFQIL
CatyCRX  DPTGTWKFTY NPMDPLDYKD QSAWKFQIL
MleuCRX  DPTGTWKFTY NPMDPLDYKD QSAWKFQIL
MmulCRX  DPTGTWKFTY NPMDPLDYKD QSAWKFQIL
MnemCRX  DPTGTWKFTY NPMDPLDYKD QSAWKFQIL
```

**Fig. S2 Alignment of primate CRX proteins.** Species name abbreviations as in Fig. 1 main text.

### Fig. S3

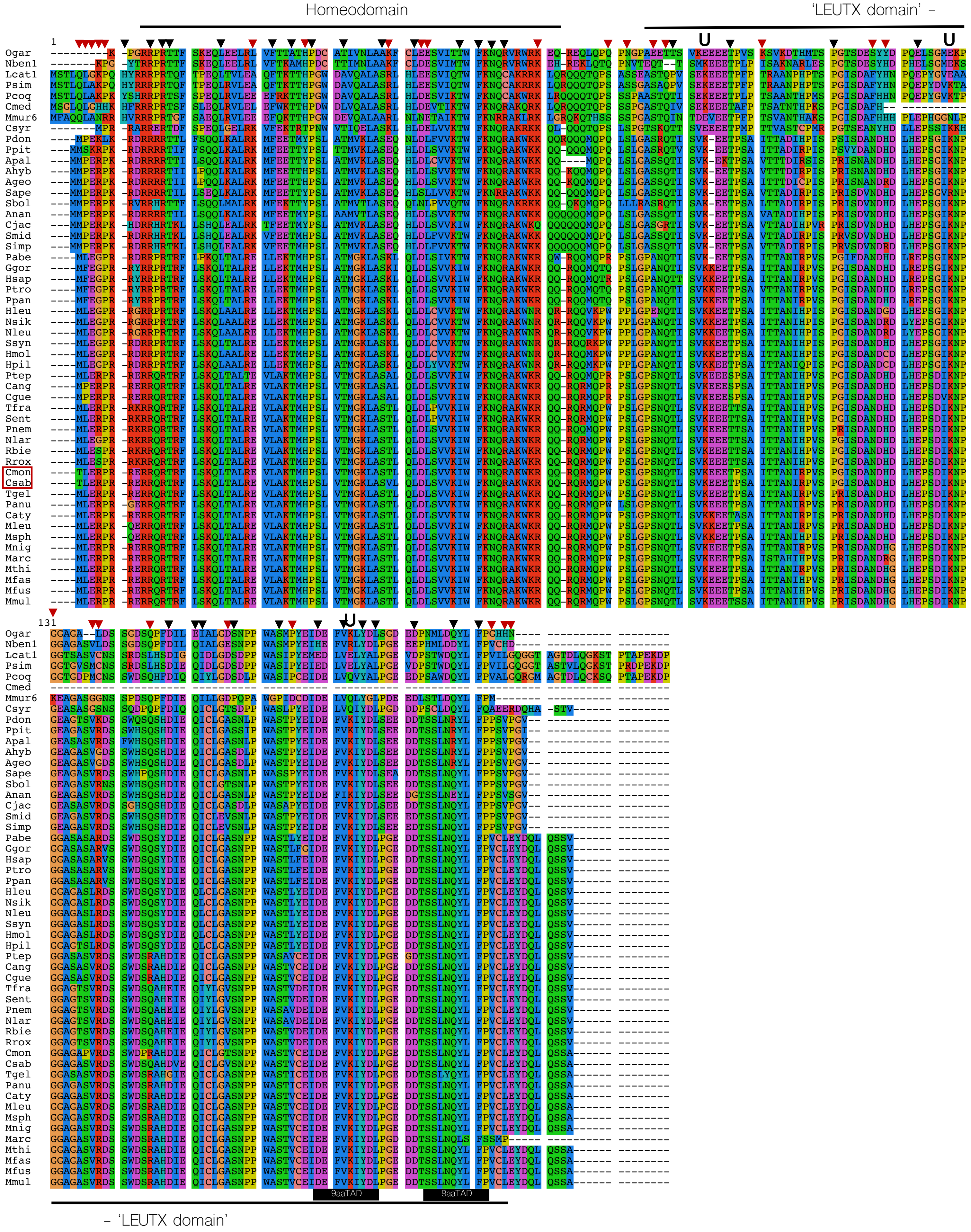

Homeodomain
'LEUTX domain' –
U
1
131
Ogar
Nben1
Lcat1
Psim
Pcoq
Cmed
Mmur6
Csyr
Pdon
Ppit
Apal
Ahyb
Ageo
Sape
Sbol
Anan
Cjac
Smid
Simp
Pabe
Ggor
Hsap
Ptro
Ppan
Hleu
Nsik
Nleu
Ssyn
Hmol
Hpil
Ptep
Cang
Cgue
Tfra
Sent
Pnem
Nlar
Rbie
Rrox
Cmon
Csab
Tgel
Panu
Caty
Mleu
Msph
Mnig
Marc
Mthi
Mfas
Mfus
Mmul
9aaTAD
9aaTAD
– 'LEUTX domain'

**Fig. S3 LEUTX protein sequence alignment used for tests for selection.** Red arrowheads indicate residues identified as under positive selection, residues marked with a black arrowhead are under pervasive purifying selection. Human ubiquitination sites are marked with a U. The position of human 9aaTADs is marked with a labelled black bar. The Cmed sequence is shortened due to an incomplete assembly, not a premature stop codon. The red square marks Cmon and Csab, in which the New World monkey start codon has mutated to a threonine residue. Ogar and Nben lack a start codon because the first LEUTX exon could not be identified due to the short nature of its length. Species name abbreviations as in Fig. 1 main text.

**Fig. S4**

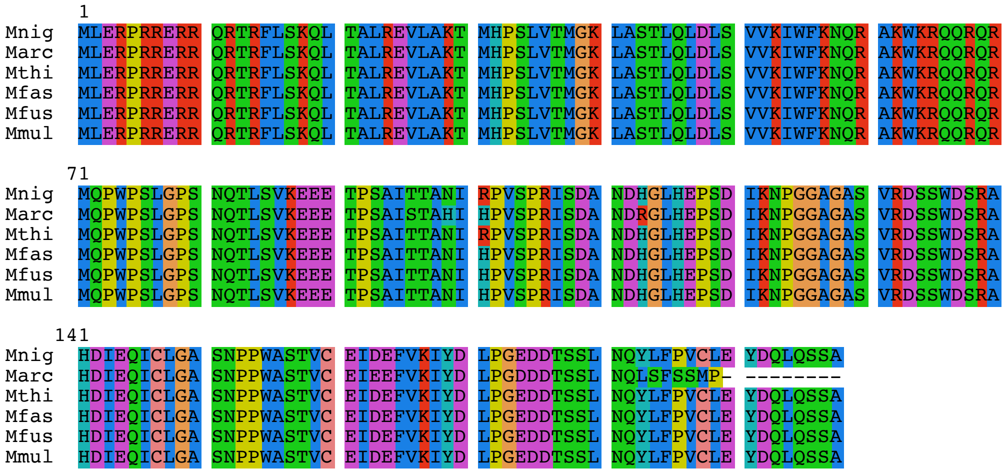


**Fig. S4 LEUTX proteins of species in the genus *Macaca*.** Abbreviations: Species abbreviations as in Fig. 1 main text.

**Fig. S5**

```
>Hsap_LEUTX_V5
ATGTTTGAGGGCCCCAGAAGATACAGACGGCCCAGAACCAGATTCCTGAGCAAGCAGCTGACAGCCCT
GAGAGAGCTGCTGGAAAAGACAATGCACCCCAGCCTGGCCACCATGGGAAAGCTGGCTTCTAAACTGC
AGCTGGACCTGAGCGTGGTCAAGATCTGGTTCAAGAACCAGCGGGCCAAGTGGAAGCGGCAGCAGAGA
CAGCAGATGCAGACCAGACCTTCTCTGGGCCCTGCCAATCAGACCACCAGCGTGAAGAAAGAGGAAAC
CCCTAGCGCCATCACCACCGCCAACATCAGACCTGTGTCTCCCGGCATCAGCGACGCCAACGATCACG
ATCTGAGAGAACCCAGCGGCATCAAGAATCCTGGCGGAGCCTCTGCCTCTGCCAGAGTGTCATCTTGG
GACAGCCAGAGCTACGACATCGAGCAGATCTGTCTGGGCGCCAGCAATCCTCCTTGGGCCAGCACACT
GTTCGAGATCGACGAGTTCGTGAAGATCTACGACCTGCCTGGCGAGGACGATACCAGCAGCCTGAACC
AGTATCTGTTCCCCGTGTGCCTGGAATACGATCAGCTGCAGAGTTCTGTTGGCGGCGGAGGATCTGGC
GGAGGCGGTTCTGGAAAGCCCATTCCTAATCCTCTGCTGGGCCTCGACAGCACCTGATGATAA


>Cjac_LEUTX_V5
ATGATGCCCGAAAGACCCAAGCACGACAGACGGCACAGAACAAAGCTGCTGAGCCACCAGCTGGAAGC
CCTGAGAAAGACCTTCGAGGAAACAATGTACCCCAGCCTGGCCACCATGGTCAAGCTGGCCTCTGAAC
AGCACCTGGACCTGAGCGTGGTCAAGATCTGGTTCAAGAACCAGCGGGCCAAGTGGAAGCAGCAGCAA
CAGCAACAACAGATGCAGCCCCAGCTGTCTCTGGGCGCCTCTGGCAGAACAATCAGCGTGAAAGAAGA
GACACCCAGCGCCGTGACCACCGCCGATATTCACCCTGTGCGGCCTAGAATCAGCGACGTGAACGACC
ACGATCTGCACGAGCCTAGCGGCATCAAGAATCCTGGCGAAGCCTCTGCCAGCGTGCGGGATTCTTCT
GGCCACAGCCAGAGCCACGACATCGAGCAGATTTGTCTGGGAGCCAGCGACCTGCCTTGGGCCTCTGC
TCCTTATGAGATCGACGAGTTCGTGAAGATCTACGACCTGTCCGAAGAGGACGACACCAGCAGCCTGA
ACCAGTACCTGTTTCCTCCAAGCGTGCCAGGCGTTGGAGGCGGAGGATCTGGCGGAGGCGGATCTGGA
AAGCCCATTCCTAATCCTCTGCTGGGCCTCGACTCCACCTGATGATAA
```

**Fig. S5 *LEUTX* expression constructs.** Sequences of ectopically expressed *LEUTX* genes, including GGGGSGGGGS linkers (blue) and V5 tag (red). Abbreviations: Cjac = *Callithrix jacchus*; Hsap = *Homo sapiens*.
